# RIPK2 dictates insulin responses to tyrosine kinase inhibitors in obese mice

**DOI:** 10.1101/2020.04.03.024620

**Authors:** Brittany M. Duggan, Joseph F. Cavallari, Kevin P. Foley, Nicole G. Barra, Jonathan D. Schertzer

## Abstract

Tyrosine kinase inhibitors (TKIs) used in cancer are also being investigated in diabetes. TKIs can improve blood glucose control in diabetic cancer patients, but the specific kinases that alter blood glucose or insulin are not clear. We sought to define the role of Receptor Interacting Serine/Threonine Kinase 2 (RIPK2) in mouse models of insulin resistance. We tested the TKI gefitinib, which inhibits RIPK2 activity, in WT, *Nod1^-/-^, Nod2^-/-^* and *Ripk2^-/-^* mice fed an obesogenic high fat diet. Gefitinib lowered blood glucose during a glucose tolerance test (GTT) in a NOD-RIPK2-independent manner in all obese mice. However, gefitinib lowered glucose-stimulated insulin secretion only in obese *Ripk2^-/-^* mice. Gefitinib had no effect on insulin secretion in obese WT, *Nodi^-/-^*, or *Nod2^-/-^* mice. Hence, genetic deletion of *Ripk2* promoted the insulin sensitizing potential of gefitinib, since this TKI lowered both blood glucose and insulin only in *Ripk2^-/-^* mice. Gefitinib did not alter the inflammatory profile of pancreas, adipose, liver or muscle tissues in obese *Ripk2^-/-^* mice compared to obese WT mice. We also tested imatinib, a TKI which does not inhibit RIPK2 activity, in obese WT mice. Imatinib lowered blood glucose during a GTT, consistent with TKIs lowering blood glucose independently of RIPK2. However, imatinib increased glucose-stimulated insulin secretion during the glucose challenge. These data show that multiple TKIs lower blood glucose, where actions of TKIs on RIPK2 dictate divergent insulin responses, independent of tissue inflammation. Our data shows that RIPK2 limits the insulin sensitizing effect of gefitinib, whereas imatinib increased insulin secretion.

## INTRODUCTION

Clinically approved tyrosine kinase inhibitors (TKIs) used in cancer are also being investigated for treatment of diabetes and diabetic complications, including retinopathy and diabetic kidney disease (1–5). However, it is still not well understood how TKIs alter blood glucose or insulin. Several TKIs lower blood glucose in humans, but the target kinases and molecular signals responsible for changes in glucose or insulin are unknown (6–15). TKIs can improve glycemic control in animal models of obesity and diabetes, but it is still not clear if inhibition of the intended kinase or off-target kinase inhibition underpins changes in glucose or insulin (16–21). Furthermore, a subset of TKIs, including nilotinib, rociletinib and ceritinib, have been associated with hyperglycemia due to impaired insulin secretion, inhibition of the insulin receptor, or development of insulin resistance (22–26). TKIs have also been reported to have opposing effects on insulin levels. Some clinical reports demonstrate increased c-peptide levels during TKI therapy (9,25) or increased β-cell insulin secretion *in vitro* (27,28), while other TKIs may impair insulin secretion (22).

Many TKIs reported to lower blood glucose share inhibitory action on molecular targets that include epidermal growth factor receptor (EGFR), c-Abl, c-Kit, and platelet-derived growth factor receptor (Pdgfrβ) (Table 1, adapted from (29)). For example, imatinib inhibits c-Abl, c-Kit, and PDGFR-β at nanomolar concentrations but imatinib has no inhibitory action on EGFR or Receptor-interacting protein kinase 2 (RIPK2). Conversely, gefitinib inhibits EGFR at a Kd value of < 1 nM and RIPK2 at a Kd value of < 530 nM. We focused on RIPK2 because this kinase propagates immune responses that have been linked to insulin resistance and dysglycemia during obesity and metabolic endotoxemia (30–32). We hypothesized that the metabolic effects of TKIs with discordant actions on RIPK2 should be tested because of the links between obesity-induced inflammation and insulin resistance. Specifically, pattern recognition receptors (PRRs) of the innate immune system have been identified as a critical point of convergence between immunity and metabolism.

**Table 1.** Inhibition of various tyrosine kinase targets by common TKIs

| **Compound** | **Tyrosine kinase targets:** |  |  |  |  |
| --- | --- | --- | --- | --- | --- |
|  | **<u>RIPK2</u>** | **<u>c-Kit</u>** | **<u>c-Abl</u>**<br>(phosphorylated) | **<u>PDGFRβ</u>** | **<u>EGFR</u>** |
| gefitinib | 530 nM | - | 480 nM | - | 1.0 nM |
| erlotinib | 680 nM | 1700 nM | 76 nM | 1400 nM | 0.7 nM |
| imatinib | - | 13 nM | 21 nM | 14 nM | - |
| sunitinib | - | 0.4 nM | 150 nM | 0.1 nM | - |
| sorafenib | 1300 nM | 28 nM | 1400 nM | 37 nM | - |
| dasatinib | 31 nM | 0.81 nM | 0.05 nM | 0.63 nM | 120 nM |
| lapatinib | 3600 nM | - | - | - | 2.4 nM |

PRRs such as Scavengers Receptors, Toll-like Receptor (TLR)s, and the Nod-like Receptor (NLR)s can propagate inflammatory signals into insulin resistance (33–37). Signals from NOD1 and NOD2 are mediated by the common downstream adaptor protein RIPK2 (38,39). We have shown that acute activation of NOD1 promotes metabolic tissue inflammation, lipolysis, insulin resistance, and dysglycemia (32). Conversely, activation of NOD2 can promote immune tolerance, lower blood glucose, and lower insulin resistance in obese mice(30–32,40). In fact, RIPK2 is required for both NOD1- and NOD2-mediated changes on blood glucose (41). It is not yet clear if drugs that target RIPK2, such as certain TKIs, allow these divergent NOD1- or NOD2-mediated responses to dominate changes in glycemia.

Several TKIs lower blood glucose and reverse insulin dependence or diabetic medication requirements in cancer patients with diabetes (6–15). TKIs that inhibit RIPK2 can influence inflammation, lipolysis, and glycemia, which are all linked in obesity. We initially hypothesized that inhibition of RIPK2 by certain TKIs (such as gefitinib) (42–44) would improve insulin sensitivity by mitigating NOD1-mediated inflammation. We also hypothesized that TKIs with no inhibitory activity towards NOD1-RIPK2 signalling would have little impact on metabolic inflammation and insulin resistance in obese mice (45). Here, we tested a TKI that inhibits RIPK2 (gefitinib) and a TKI that does not inhibit RIPK2 (imatinib) in obese mice.

In contrast to our initial hypothesis, we found that RIPK2 mediated the effects of TKIs on insulin rather than glucose. We found that deletion of RIPK2 improved the insulin sensitizing potential of gefitinib. We found that gefitinib lowered oral glucose-stimulated insulin secretion in obese *Ripk2^-/-^*, but not in obese WT, *Nod1^-/-^* or *Nod2^-/-^* mice, while blood glucose was lowered during glucose challenge in all genotypes of mice. We also found that imatinib, a TKI that does not inhibit RIPK2, increased insulin secretion. Our data show that TKIs can lower blood glucose independently of NOD-RIPK2 signalling. However, inhibition of RIPK2 signalling by certain TKIs and actions on insulin secretion should be carefully considered in patients and before retasking these drugs for diabetes, its complications or aspects of metabolic disease. Importantly, the various target kinases acted upon by TKIs must be considered in discordant glucose and insulin responses, since imatinib lowered glucose, but increased insulin levels, during a glucose load in obese mice.

## MATERIALS AND METHODS

### Mice and materials

All procedures followed the Canadian Council on Animal Care (CCAC) and approved by McMaster University Animal Ethics Review Board (AREB). Genotyping analysis based on the nucleotide transhydrogenase (*Nnt*) gene was used to identify the substrains of our knockout models (*Nod1^-/-^, Nod2^-/-^* and *Ripk2^-/-^* mice). Mice used in these studies were typical of the C57BL/6J (WT/J) or C57BL/6N (WT/N) background. We did not compare results between substrains or different knockout mice. Our goal was to test the effects of TKIs, hence we tested drug treatment in both HFD-fed WT/J and WT/N mice to rule out any effect of the background strain on TKI-mediated glucose and insulin effects. WT/J, *Nod1^-/-^, Nod2^-/-^* and *Ripk2^-/-^* mice were born under specific pathogen-free conditions at McMaster University. WT/N mice were purchased from Taconic Biosciences (Model# DIO-B6-M; Rensselaer, NY). Male mice were used. Gefitinib was from AbMole Bioscience (Houston, TX). Imatinib was from AdooQ Bioscience (Irvine, CA). Methylcellulose (M0512) used as a vehicle for TKI oral delivery was from Sigma-Aldrich (St. Louis, MO). Muramyl dipeptide (MDP) (Cat# tlrl-mdp) and ultrapure LPS from *Escherichia coli O111:B4* (Cat# tlrl-3pelps) were from InvivoGen (San Diego, CA). Blood glucose was repeatedly measured via tail vein sampling using Accu-Chek Aviva blood glucometer from Roche Diagnostics (Missisauga, ON).

#### Animal protocols

All mice were maintained under controlled lighting (12:12 Light:Dark) and temperature (22°C) with *ad libitum* access to food and water. TKIs were suspended in 1% methylcellulose at 10-50mg/mL with brief sonication followed by vortexing to achieve a homogenous colloid. Mice were administered 150-300μL via oral gavage based on individual body weight to achieve the reported dose (in mg/kg).

##### Obesity models

All mice were 8-10 weeks of age before being fed an obesogenic diet containing 60% kcal from fat (Research Diets, D12492). Where indicated, age-matched mice were also fed a control diet (Teklad 22/5, 17% kcal from fat). After 10 weeks of high fat diet (HFD) feeding, gefitinib (50 mg/kg) or imatinib (250 mg/kg) was orally administered every second day for 8 treatments. Then, a 6h-fasted glucose tolerance (GTT) or insulin tolerance test (ITT) was performed (HFD; 1g/kg, i.p., Control diet; 2g/kg, i.p.). Body fat composition was measured using whole body MRI (Bruker Minispec, LF90-II) after the 10^th^ TKI treatment. Mice were provided 11 total treatments over 22 days before assessing oral glucose-stimulated insulin secretion (OGSIS) as detailed in Figure 1. Blood samples were collected via tail-vein sampling at t=0, 10 and 60m post-glucose gavage (4g/kg). Blood was centrifuged for 10 min at 4°C and 10,000 g, and plasma fraction was collected and stored at −80°C. Plasma insulin was assessed using high sensitivity mouse insulin ELISA kit (Toronto Bioscience, Cat# 32270) and measured with a Synergy H4 Hybrid reader (Biotek Instruments). Area under the curve (AUC) was calculated for GTT, ITT and OGSIS results using GraphPad Prism 6 software. HOMA-IR was calculated by multiplying blood glucose values (mmol/L) by insulin values (μU/mL) and dividing this by 22.5, as described (46). Insulin resistance index was calculated by multiplying GTT AUC by OGSIS AUC by 10^-4^, as described (47). Following OGSIS, mice were euthanized by cervical dislocation and liver, epidydimal adipose, pancreas and tibialis anterior muscles were rapidly excised, weighed and snap frozen in liquid nitrogen.

**Figure 1.**
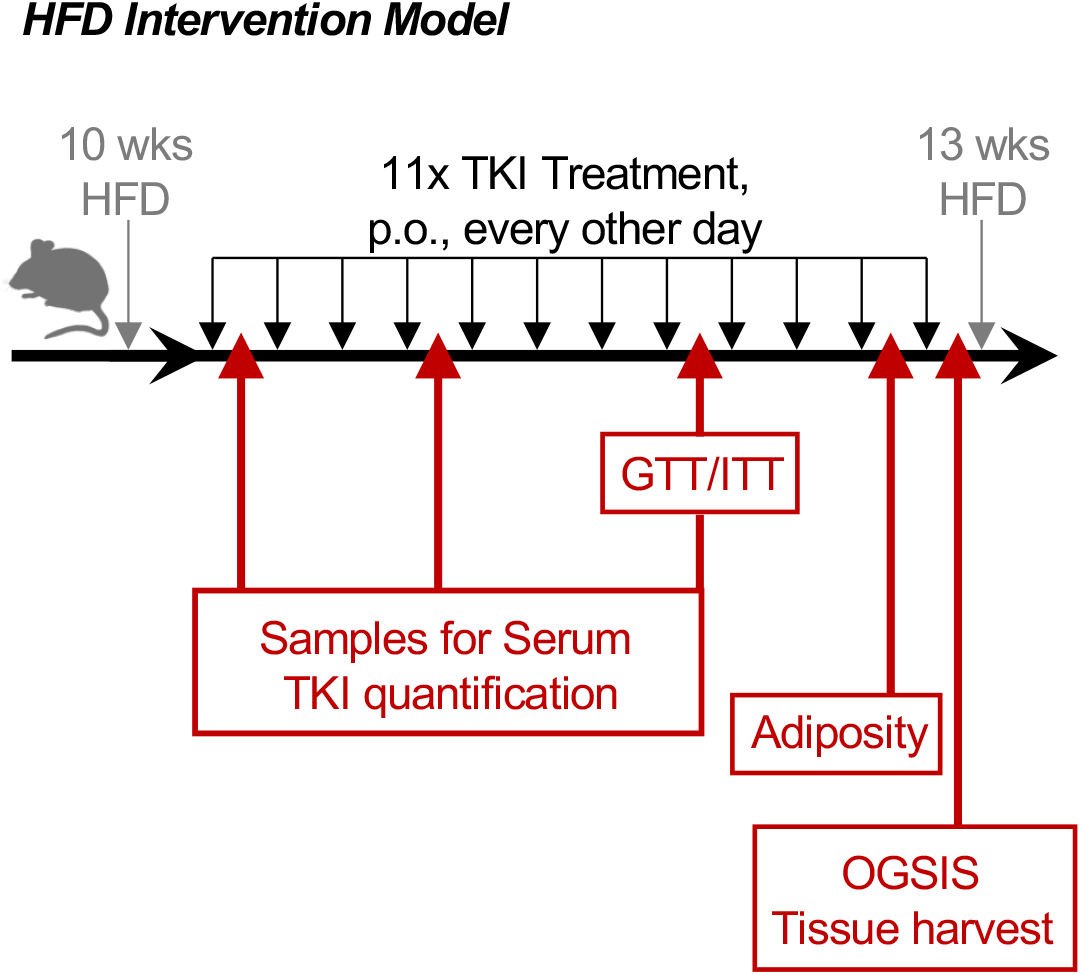
Experimental design for TKI administration and testing glucose and insulin dynamics in obese mice. Mice were fed a 60% HFD for 10 weeks before intervention with tyrosine kinase inhibitor (TKI, p.o., every other day). The TKIs used were gefitinib (50 m/kg) or imatinib (250 mg/kg). Mice received 8x treatments over 16 days before GTT or ITT was assessed. Blood samples were collected after the 1st, 4th and 8th treatment for quantification of TKI concentration in serum. Adiposity was assessed after 10 treatments and oral-glucose stimulated insulin secretion (OGSIS) was assessed after 11 treatments. Metabolic tissues were harvested immediately following cervical dislocation after blood samples were collected for measuring insulin. GTT, glucose tolerance test; ITT, insulin tolerance test; OGSIS, oral glucose-stimulated insulin secretion.

##### Acute NOD2 activation model

WT/J mice were orally administered equimolar doses of gefitinib (100mg/kg, p.o.), imatinib (110mg/kg) or methylcellulose vehicle control and then 4 hours later were injected with MDP (100μg, i.p) or saline vehicle, each day for 3 days. On the 4^th^ day, all mice were injected with ultrapure LPS (0.2mg/kg, i.p.), fasted for 6h, and a GTT was performed (2g/kg, i.p.).

### Gene expressional analysis

Total RNA was obtained from ~50 mg of liver, epidydimal adipose and tibialis anterior muscle via mechanical homogenization at 4.5 m/s for 30 s using a FastPrep-24 tissue homogenizer (MP Biomedicals) and glass beads, followed by phenol-chloroform extraction. RNA was treated with DNase I (Thermo Fisher Scientific) and cDNA was prepared using 500-1000 ng total RNA and SuperScript III Reverse Transcriptase (Thermo Fisher Scientific). Transcript expression was measured using TaqMan Assays with AmpliTaq Gold DNA polymerase (Thermo Fisher Scientific) and target genes were compared to the mean of *Rplp0* and *18S* housekeeping genes using the ΔΔ*C*_T_ method. A list of target probe sequences can be found in the online archive containing supplemental data (48).

### Quantification of serum TKI concentration

#### Quantification of serum TKI concentration after single administration in control diet-fed mice

WT/J mice were orally administered an equimolar dose of gefitinib (100mg/kg, p.o.), imatinib (110mg/kg) or methylcellulose vehicle control. These doses correspond to the doses used in control diet-fed mice during acute NOD2 activation experiments. Blood samples were collected via tail vein sampling at t=0, 2 and 6h post-administration.

#### Quantification of serum TKI concentration during repeated administration in obese mice

Obese WT/J mice fed the HFD for 10 weeks were orally administered gefitinib (50mg/kg, p.o.), imatinib (250mg/kg) or methylcellulose (vehicle control). These doses correspond to the doses used in obese mice in our HFD intervention model. Blood samples were collected via tail vein sampling 2h after the 1^st^, 4^th^ and 8^th^ administration (on treatment days 1, 7, and 15).

#### TKI serum sample processing and quantification

All blood samples were clotted at room temperature for 20 min, then centrifuged for 10 min (4°C at 10,000 g) and serum fraction was collected and stored at −80°C until analysis. Quantification was accomplished using liquid chromatography coupled to tandem mass spectrometry based on a previously published method (49). In brief, an acetonitrile precipitation was performed prior to sample analysis by an Agilent 1290 Infinity II HPLC with an Agilent 6550 iFunnel Q-TOF mass spectrometer for detection (Agilent, Santa Clara, CA, USA). Gefitinib and deuterated gefitinib (gefitinib-d6, Cayman Chemicals, Ann Arbor, MI), imatinib and deuterated imatinib (imatinib-d3, Cayman Chemicals, Ann Arbour, MI) were used to generate a matrix-matched standard curve and determine sample recovery. Method detection was confirmed in a linear concentration range, reproducibility was verified, matrix effects were tested, and three TKI concentrations were validated (0.1ppm-20ppm). No gefitinib or imatinib was detected in control samples above the method detection limit of 0.02ppm.

### Data analysis

Data is expressed as mean ± standard error of the mean (SEM). Comparisons were made using unpaired, two-tailed Student’s t-test, where 2 variables are compared. ANOVA was used for comparison of more than 2 variables and Tukey’s post hoc test was used when appropriate (Prism 4-6; Graphpad Software). Differences were considered statistically significant at p <0.05.

## RESULTS

### RIPK2 limits insulin secretion and the insulin-sensitizing potential of the TKI gefitinib

Mouse substrain notation is critical in genetically modified mouse models (50–52). We genotyped all mice for heterogeneity in the nucleotide transhydrogenase (*Nnt*) gene to identify the substrains of our knockout models (*Nod1^-/-^, Nod2^-/-^* and *Ripk2^-/-^* mice) used in these studies and found that all of our models either existed on either C57BL/6J or C57BL/6N background. To reconcile any influence of the substrain background or *Nnt* expression on the effect of TKIs, we first assessed the effect of gefitinib treatment (50mg/kg, p.o) on glucose levels during a GTT and OGSIS in HFD-fed WT/J and C57BL6/N (WT/N) mice, in addition to HFD-fed *Nod1^-/-^, Nod2^-/-^* and *Ripk2^-/-^* mice (Figure 1).

We measured insulin secretion and the Insulin Resistance Index (47) during fasting and during an oral glucose load (4g/kg, D-glucose, p.o) in mice treated with vehicle or gefitinib (50mg/kg, p.o., every other day). Gefitinib did not change fasting blood glucose or HOMA-IR in obese WT/J, WT/N, *Nod1^-/-^* or *Nod2^-/-^* or *Ripk2^-/-^* mice (Figure 2A-E). Gefitinib did not change glucose-stimulated blood insulin levels and did not alter the Insulin Resistance Index in HFD-fed, obese WT/J, WT/N, *Nod1^-/-^* or *Nod2^-/-^* mice (Figure 3A-D). However, gefitinib lowered glucose-stimulated insulin secretion and lowered the Insulin Resistance Index in obese, HFD-fed *Ripk2^-/-^* mice (Figure 3E).

**Figure 2.**
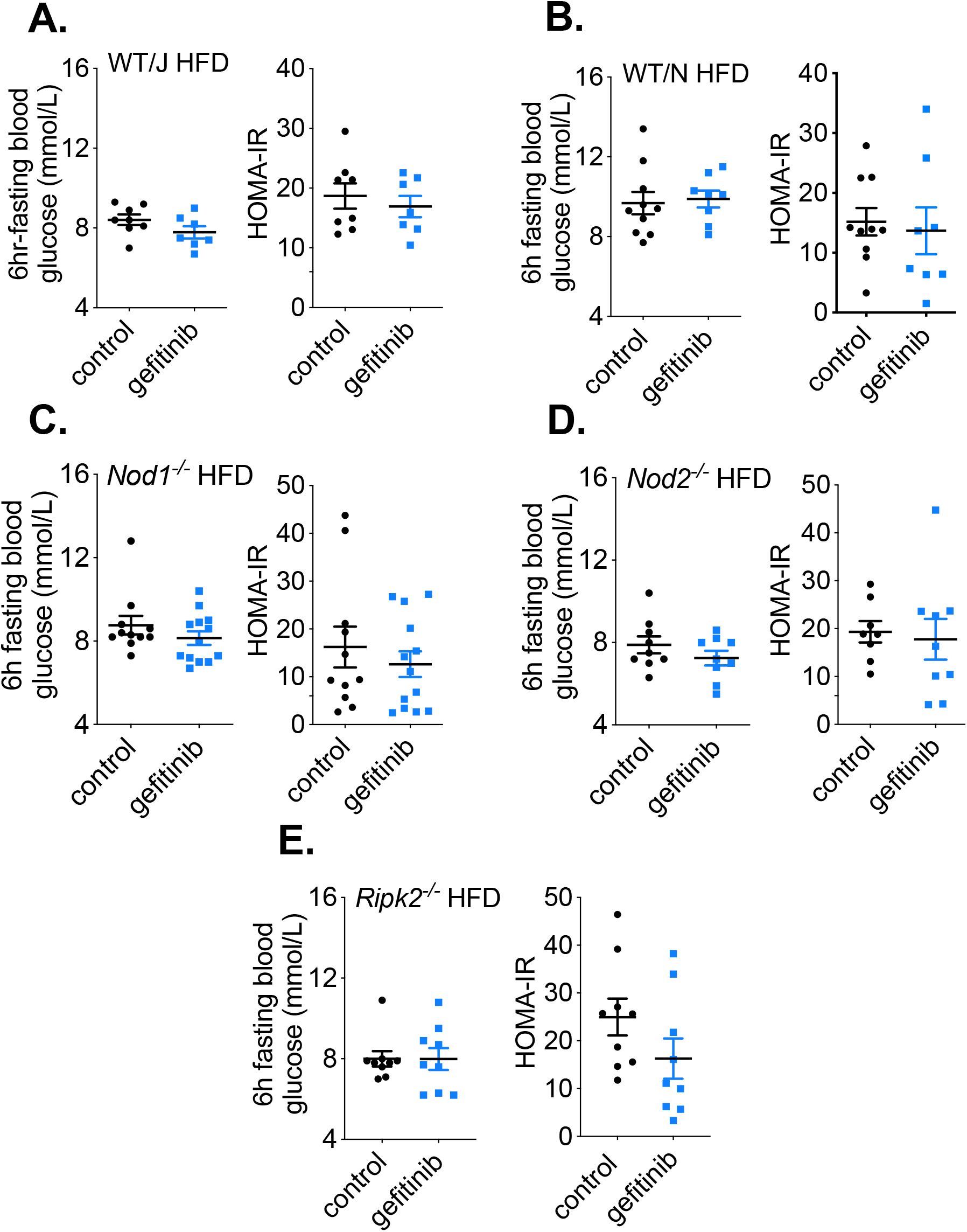
Gefitinib does not alter fasting glycemia or HOMA-IR in obese mice. 6 h fasting blood glucose and HOMA-IR was calculated after 11 treatments of gefitinib (50mg/kg, or vehicle, p.o., every other day) in A) WT/J mice, B) WT/N mice, C) *Nod1^-/-^* mice, D) *Nod2^-/-^* mice and E) *Ripk2^-/-^* mice fed a HFD for 10wks before initiating gefitinib treatment. Values are mean ± SEM. *Denotes statistical differences between groups (p<0.05). Each dot indicates a mouse.

**Figure 3.**
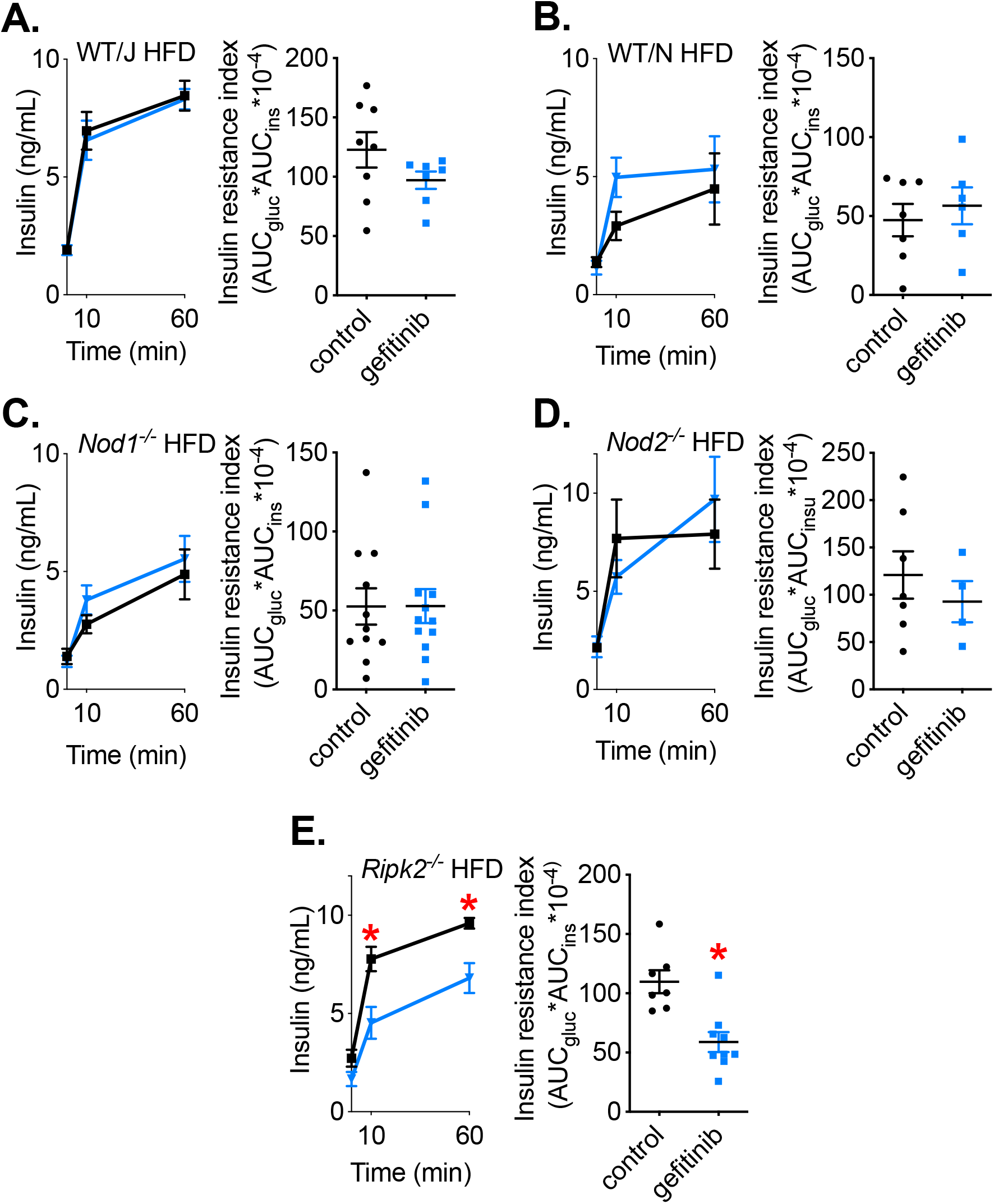
Deletion of RIPK2 lowers both insulin and glucose and improves the insulin sensitizing potential of gefitinib. Blood insulin levels before and t=10m, 60m following an oral glucose challenge (4g/kg, p.o.) after 11 treatments of gefitinib (50mg/kg, or vehicle, p.o., every other day) and Insulin Resistance Index during the glucose load was calculated in HFD-fed A) WT/J, B) WT/N, C) *Nod1^-/-^*, D) *Nod2*^-/-^ and E) *Ripk2^-/-^* mice. Values are mean ± SEM. *Denotes statistical differences between groups (p<0.05). Each dot indicates a mouse.

### Gefitinib improves glucose control in obese mice without changing body weight or adiposity

Glucose tolerance tests were conducted after 8 treatments of gefitinib in obese WT/J and WT/N mice that were fed an obesogenic HFD for 10 weeks prior to initiating TKI treatment. Gefitinib lowered glucose during a GTT in both obese WT/J and WT/N mice (Figure 4A-B). We also tested a cohort of age-matched, control diet-fed WT/J mice to determine if glucose tolerance was altered by gefitinib treatment in lean mice. In contrast to obese mice, gefitinib treatment did not alter glucose tolerance in lean mice (Figure 4C). This is consistent with a report showing that the TKI imatinib does not alter glucose control in lean mice, but improves glucose control in obese *db/db* mice (16). We also found that gefitinib treatment promoted lower blood glucose during an ITT in HFD-fed WT/J mice (Figure 4D). Importantly, the glucose lowering effects of gefitinib occurred independently of any changes in body mass or adiposity in WT/J mice (Figure 4E-F). Gefitinib treatment did not alter food consumption at any timepoint during the treatment period in mice (data not shown).

**Figure 4.**
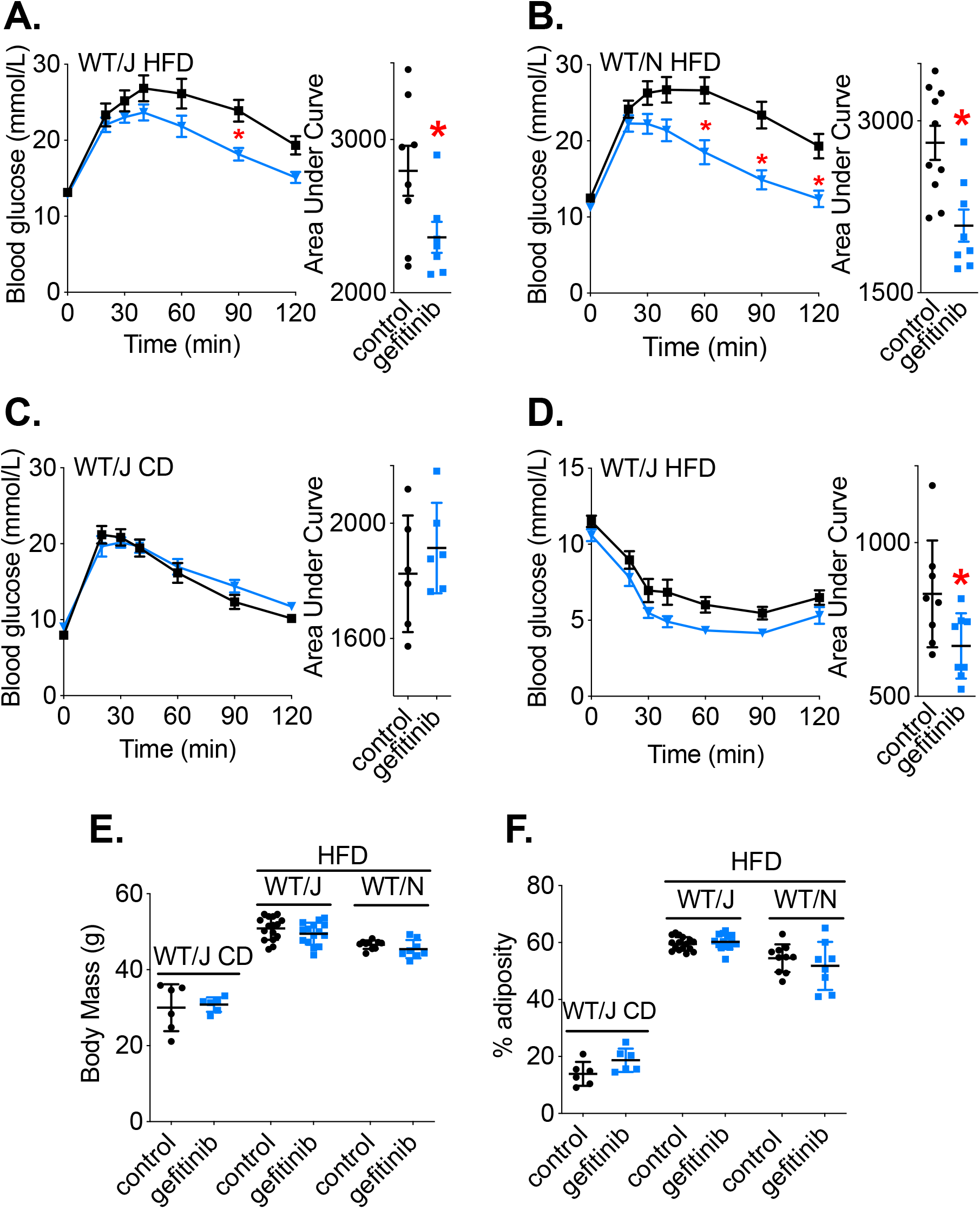
Gefitinib attenuates impaired glucose control and insulin resistance in obese mice independent of changes in body mass or adiposity. Blood glucose vs. time and area under the curve (AUC) after 8 treatments of gefitinib (50mg/kg, or vehicle, p.o., every other day) during a glucose tolerance test (1g/kg, i.p.), in A) WT/J or B) WT/N mice fed a HFD for 10 weeks before initiating gefitinib treatment, or C) in age-matched lean WT/J mice fed a control diet (GTT: 2g/kg, i.p.). D) Blood glucose vs. time and AUC after 8 treatments of gefitinib (50mg/kg, or vehicle, p.o., every other day) during an insulin tolerance test (2 IU/kg, i.p.) in obese WT/J mice fed HFD for 10 weeks before initiating gefitinib treatment. E) Body mass and F) percent adiposity of all control-fed WT/J, and HFD-fed WT/J and WT/N mice. Values are mean ± SEM. *Denotes statistical differences between groups (p<0.05). Each dot indicates a mouse.

### Gefitinib improves glucose control, independently of *Nod1^-/-^, Nod2^-/-^* or *Ripk2^-/-^*

We also assessed gefitinib treatment (50mg/kg, p.o., every other day) in obese *Nod1^-/-^, Nod2^-/-^* and *Ripk2^-/-^* mice. Compared to vehicle treated mice, gefitinib lowered blood glucose at specific times during a GTT in all mouse genotypes (Figure 5A-C). Gefitinib treatment did not alter body mass or adiposity in *Nod1^-/-^, Nod2^-/-^* and *Ripk2^-/-^* mice (Figure 5D-E). Given that *Ripk2^-/-^* mice treated with gefitinib require less insulin to achieve lower blood glucose levels during a glucose load (Figure 3E), and the lower Insulin Resistance Index only in *Ripk2^-/-^* mice treated with gefitinib (Figure 3E), these data support the conclusion that deletion of RIPK2 increases the insulin-sensitizing potential gefitinib. Our data points to an effect of RIPK2 on gefitinib-mediated effects during a glucose load rather than in the fasting state.

**Figure 5.**
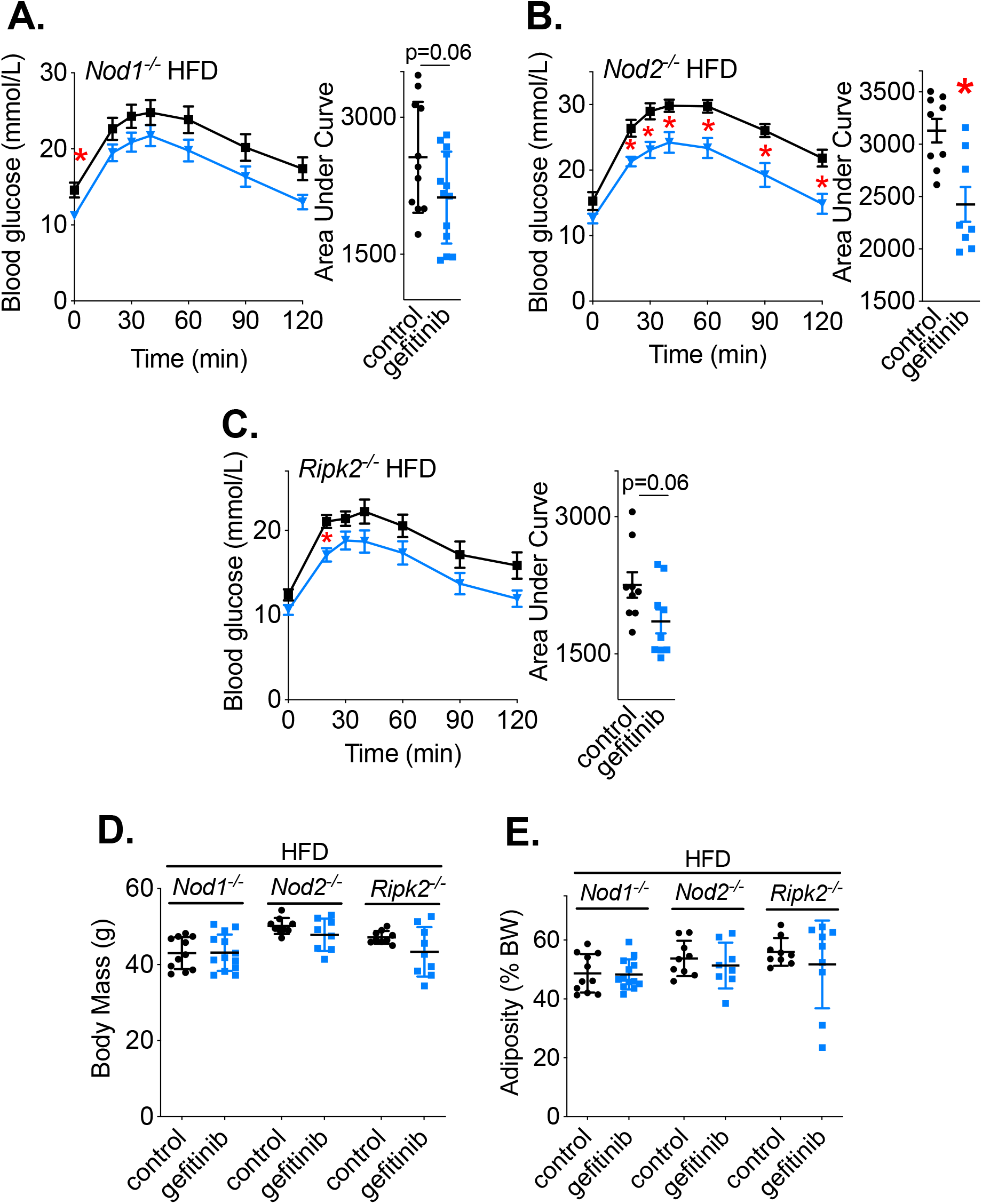
Gefitinib attenuates impaired glucose control in obese mice independent of NOD1, NOD2 or RIPK2. Blood glucose vs. time and calculated area under the curve (AUC) after 8 treatments of gefitinib (50mg/kg, or vehicle, p.o., every other day) during a glucose tolerance test (1g/kg, i.p.), in A) *Nod1^-/-^*, B) *Nod2^-/-^* or C) *Ripk2^-/-^* mice fed a HFD for 10 weeks before initiating gefitinib treatment. D) Body mass and E) percent adiposity of HFD-fed *Nod1^-/-^, Nod2^-/-^*, and *Ripk2^-/-^* tested. Values are mean ± SEM. *Denotes statistical differences between groups (p<0.05). Each dot indicates a mouse.

### Gefitinib does not cause widespread changes in markers of inflammation or ER stress in metabolic tissues

At least two previous reports demonstrated decreased inflammation in liver, adipose and muscle, or decreased ER stress in liver and adipose, of obese mice treated with TKIs (16,18). Thus, we next measured gene expression of a panel of inflammatory and ER stress-related genes in liver, adipose, and muscle. To address the role of inflammation and ER stress on divergence in insulin responses between WT/J vs. *Ripk2^-/-^* mice, we assessed gene expression in WT/J and *Ripk2^-/-^* mice to see if there was a divergence in inflammatory or ER stress-related genes that paralleled the metabolic differences.

Lower expression of *cxcl1* in liver was observed in WT/J and *Ripk2^-/-^* mice treated with gefitinib, and lower expression of *Tnf* in liver and lower *Cxcl1* in muscle was observed in WT/J mice treated with gefitinib. The reader is referred to the online archive for all transcript data in liver, muscle and adipose tissues (48). Overall, there were no widespread differences in pro-inflammatory cytokine gene expression in liver, muscle, or adipose tissue of gefitinib-treated mice and no robust pattern of changes occurred due to gefitinib treatment between WT/J versus *Ripk2^-/-^* mice. Changes in expression of some anti-inflammatory markers (*irak3, tgfb1*) were observed in adipose and muscle of WT/J or *Ripk2^-/-^* mice. However, no consistent or robust increases in anti-inflammatory markers were observed in either WT/J or *Ripk2^-/-^* mice. Similarly, no significant changes in gene expression of immune cell markers were observed in any tissue of WT/J or *Ripk2^-/-^* mice. Finally, a decrease in the spliced form of *xbp1 (sxbp1)* was observed in liver of *Ripk2^-/-^* mice. In conclusion, changes in a limited number of inflammatory or ER stress-related genes in metabolic tissues did not correspond with the effects of gefitinib on insulin secretion or sensitivity. These inflammatory markers in adipose, liver and muscle are not positioned to account for the robust improvements in glucose tolerance observed with TKI treatment in diet-induced obese mice or the RIPK2-mediated changes in insulin secretion during a glucose load.

### TKIs that inhibit RIPK2 prevent the insulin-sensitizing properties of postbiotics

It was known that gefitinib prevents dysglycemia caused by acute NOD1 activation (45). It was also known that RIPK2 in non-hematopoietic cells is required for the NOD2-activating ligand MDP to improve blood glucose control (40,41). RIPK2 is the adapter for both NOD1 and NOD2 and here we sought to test if only TKIs that block RIPK2 would prevent MDP/NOD2-mediated changes in blood glucose. We used our established model of acute NOD2-activation with the insulin sensitizer MDP during endotoxin challenge to test a TKI that inhibits RIPK2 (gefitinib) versus a TKI that does not affect RIPK2 (imatinib) (29,45). As expected, 3 days of MDP treatment (100μg, i.p.) lowered fasting blood glucose and improved glucose tolerance during an endotoxin challenge (LPS, 0.2mg/kg, i.p.) without altering body mass (Figure 6A-C). Gefitinib treatment (100mg/kg/day, p.o) prevented MDP-induced lowering of blood glucose in the fasted state and during the GTT (Figure 6B-C). In contrast, treatment with an equimolar dose of imatinib (110mg/kg/day, p.o.) did not block MDP-lowering of fasting blood glucose or lower blood glucose during the GTT during an endotoxin challenge (Figure 6E-G).

**Figure 6.**
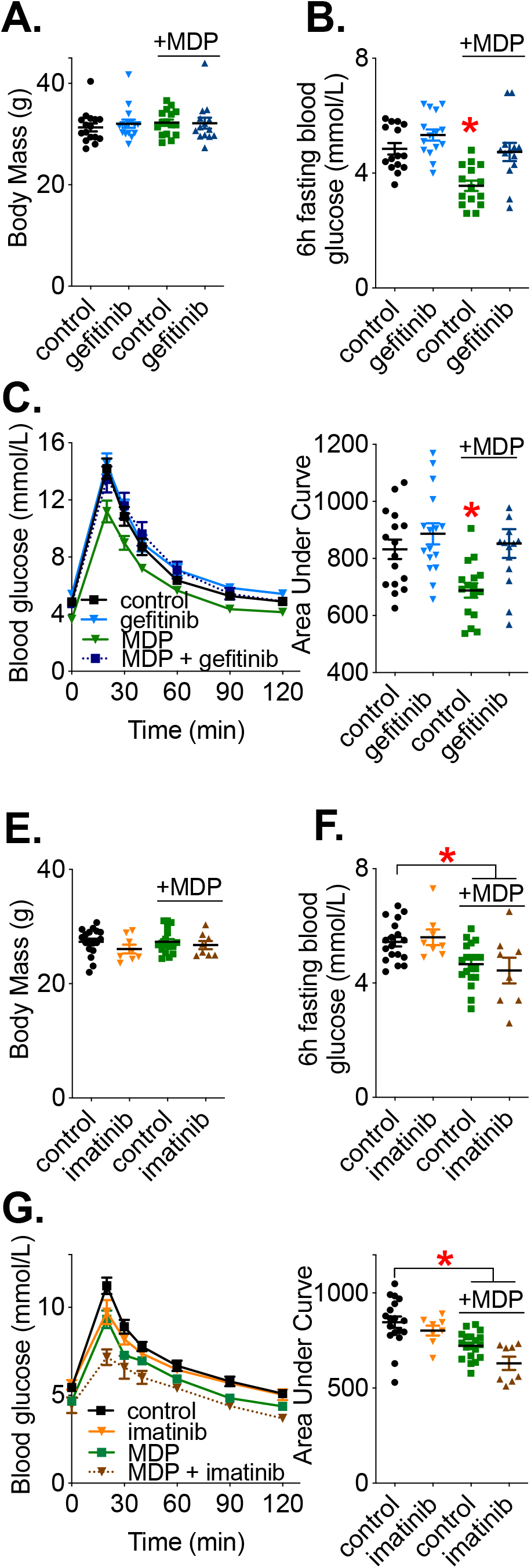
The TKI gefitinib, but not imatinib, attenuates NOD2-mediated improvements in glucose control during mild endotoxin challenge in mice. A) Body mass, B) 6h fasting blood glucose and C) Blood glucose vs. time and AUC in mice treated with gefitinib, 100 mg/kg p.o., or vehicle, and injected with MDP (100μg, i.p.), for 3 days prior to induction of mild, acute endotoxemia (0.2 mg/kg LPS, i.p.) on the 4th day, then a GTT was performed 6h post-LPS injection. D) Body mass, E) 6 h fasting blood glucose and F) Blood glucose vs. time and AUC in mice treated with imatinib (110 mg/kg p.o, or vehicle, and injected with MDP (100 μg, i.p.), for 3 days prior to induction of endotoxemia (0.2mg/kg LPS, i.p.) on the 4th day, then a glucose tolerance test was performed 6h post-LPS injection. Values are mean ± SEM. *Denotes statistical differences between groups (p<0.05). Each dot indicates a mouse.

### Imatinib lowers blood glucose, but increases insulin during a glucose load in obese mice

We next sought to test how imatinib, a TKI that does not inhibit RIPK2, influences insulin and glucose in obese, HFD-fed mice. Imatinib has no reported inhibitory effects against RIPK2 (29) and our published data demonstrates that imatinib does not inhibit the acute metabolic or immune effects of NOD1 activation (45). Additionally, above we show that imatinib does not inhibit the acute effects of NOD2 activation on glycemia (Figure 6E-G). Thus, we initially hypothesized that imatinib treatment would lower both blood glucose and insulin levels in obese mice, mimicking our findings in *Ripk2^-/-^* mice treated with gefitinib. Initially, we tested an equimolar dose of imatinib (55mg/kg, p.o., every other day) as used in our gefitinib HFD intervention model in WT/J mice. However, this dose of imatinib did not significantly improve blood glucose or alter insulin secretion during a glucose challenge (data not shown). We then established that a higher dose of imatinib, 250mg/kg, p.o., could be administered for the duration of the treatment period without altering body mass or adiposity (Figure 7A-B). When using this higher dose of imatinib (250mg/kg, p.o.), we found that imatinib-treated mice had significantly lower glucose during a GTT (Figure 7C). However, contrary to our initial hypothesis, we found that imatinib-treated obese mice had increased HOMA-IR and higher glucose-stimulated blood insulin levels compared to vehicle-treated mice (Figure 7E-F). The increase in insulin secretion coupled with lowering of blood glucose equated to no change in the Insulin Resistance Index due to imatinib treatment of obese mice (Figure 7G). This data supports a model where various TKIs achieve their effects on blood glucose regulation via different mechanisms including higher insulin secretion (imatinib) or increased insulin sensitivity (gefitinib).

**Figure 7.**
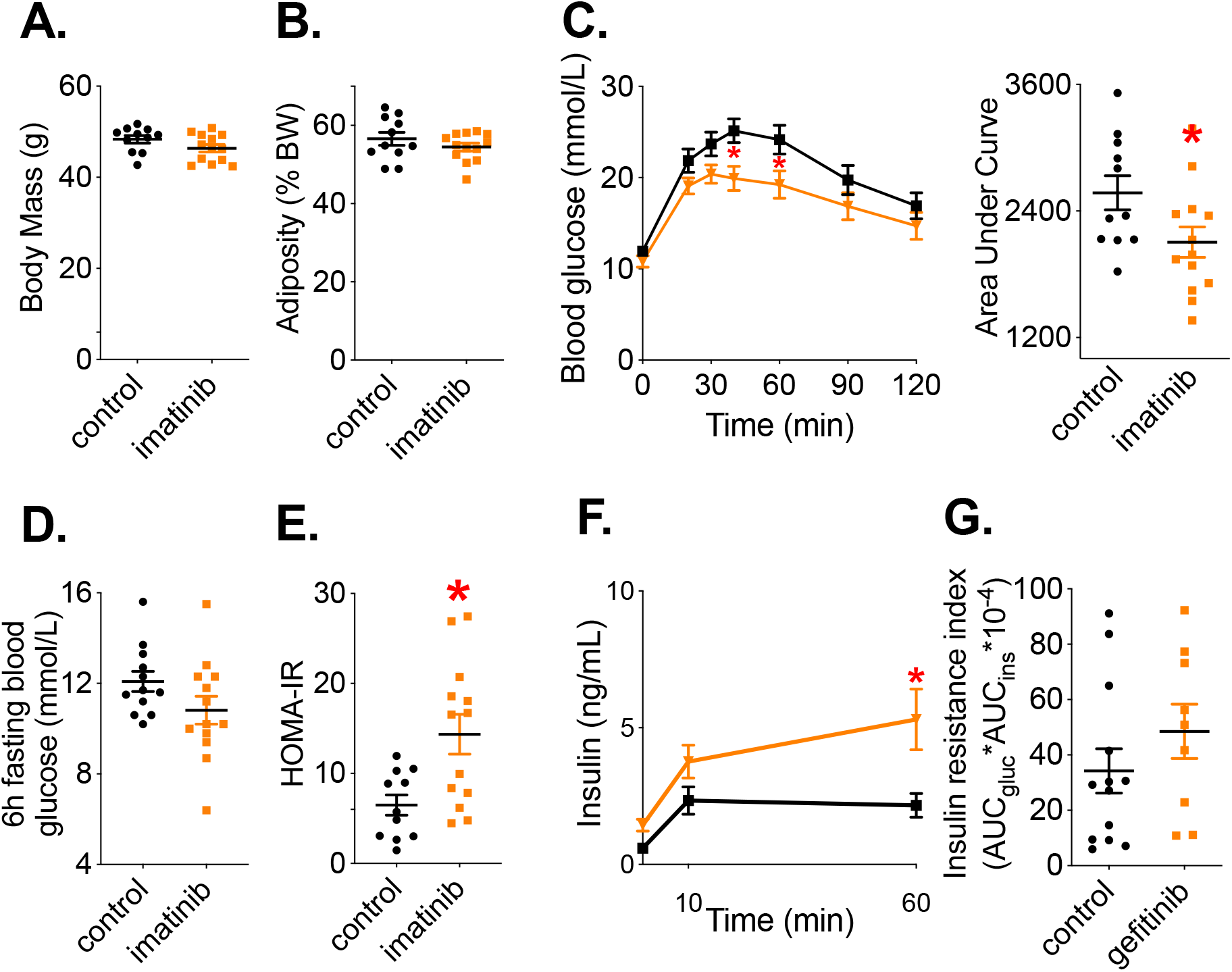
Imatinib improves glucose tolerance but induces hyperinsulinemia during obesity. A) Body mass and B) percentage adiposity (B) after 8 treatments of imatinib (250mg/kg, or vehicle, p.o., every other day) in WT/J mice fed an obesogenic HFD for 10 weeks before initiating treatment regime. Blood glucose vs. time (C) and AUC (inset) during a 6h fasted glucose tolerance test. Blood insulin levels before and at t=10, 60 min following an oral glucose challenge (4g/kg, p.o.) after 11 treatments of imatinib (D) and Insulin Resistance Index was calculated (E). Values are mean ± SEM. * denotes statistical differences between groups (p<0.05). Each dot indicates a mouse.

### TKIs do not cause widespread changes in pancreatic expression of inflammatory or ER-stress markers

Our data shows that obese *Ripk2^-/-^* mice treated with gefitinib secrete less insulin during a glucose challenge, whereas insulin secretion is not altered in obese WT/J mice treated with gefitinib. In contrast, obese WT/J mice treated with imatinib exhibit increased insulin secretion during a glucose challenge. Considering the divergent effects on insulin secretion in WT/J vs. *Ripk2^-/-^* mice, and with gefitinib vs. imatinib, we next assessed gene expression of a panel of inflammatory and ER stress-related genes in pancreas tissue of WT/J and *Ripk2^-/-^* mice treated with gefitinib, and in WT/J mice treated with imatinib. The goal was to assess if there was a pancreas-specific immune or ER stress effect that could be driving divergence in insulin secretion caused by administration of these TKIs.

Decreased expression of *cxcl1* and *il1b* in pancreas was observed in both *WT/J* and *Ripk2^-/-^* mice treated with gefitinib (Figure 8A-B, top left). Gefitinib did lower the expression of a small number of anti-inflammatory markers in pancreas such as *irak3* in WT/J mice and *arg1* in *Ripk2^-/-^* mice (Figure 8A-B, top right). No significant changes in gene expression of immune cell markers or ER stress-related genes were observed (Figure 8A-B, bottom). In WT/J mice treated with imatinib, we found significantly lower levels of *cxcl10*, and significantly increased levels of *il4* in the pancreas (Figure 8C). Overall, no robust or consistent pattern of changes in inflammatory- or ER stress-related genes were observed in pancreas tissue from WT/J or *Ripk2^-/-^* mice treated with gefitinib or WT/J mice treated with imatinib that could account for RIPK2-mediated divergence in glucose-stimulated insulin secretion.

**Figure 8.**
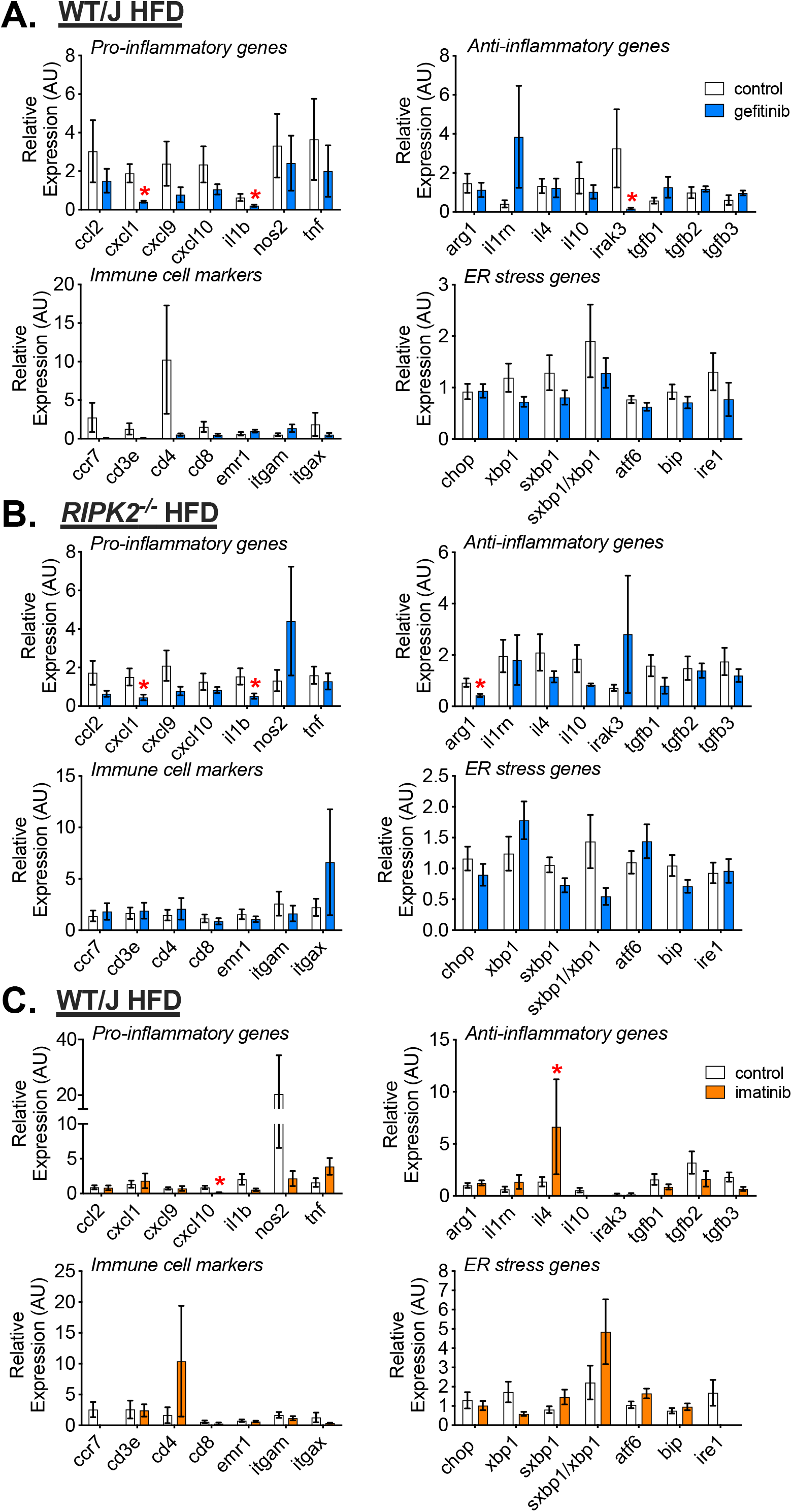
The TKIs gefitinib and imatinib do not cause widespread changes in inflammatory or ER-stress related gene profiles in pancreas of obese mice. Transcript levels of pro-inflammatory cytokines (top left), anti-inflammatory cytokines (top right), immune cell markers (bottom left) and ER-stress related genes (bottom right) in pancreas tissue of A) obese WT/J treated with gefitinib, B) obese *Ripk2^-/-^* mice treated with gefitinib, and C) obese WT/J mice treated with imatinib. Values are mean ± SEM. * denotes statistical differences between groups (p<0.05). n = 3-9 per group.

### Gefitinib and imatinib dosing resulted in comparable serum drug concentrations

We quantified serum concentrations of each TKI throughout our treatment period to validate the doses of gefitinib (50mg/kg) and imatinib (250mg/kg) used in obese mice. These were the doses required for each TKI to improve glucose tolerance in obese mice without weight loss or an anorexic effect. The pharmacokinetic and pharmacodynamic properties of gefitinib and imatinib have been well studied in lean mice, but to the best of our knowledge, no published report in the literature exists of blood concentrations following TKI gefitinib and imatinib administration to obese mice. Furthermore, it was important to understand how TKIs could accumulate with repeated administration, as in our HFD-fed mouse model, given the different oral doses used to achieve lowered blood glucose. First, we sought to quantify serum TKI concentration after a single TKI administration in lean, control diet-fed mice to compare to other published results in the literature. Mice were given a single administration of gefitinib (100mg/kg, p.o.) or an equimolar dose of imatinib (110mg/kg, p.o.), that correspond to doses used in acute NOD2 experiments in lean mice (see Figure 6). We have also shown this dose of gefitinib effectively inhibits acute NOD1 signalling *in vivo* (45) and these doses are comparable to what is used in many published mouse xenograph models (53–58). Gefitinib serum concentrations were 3.857 ± 0.263 μg/mL and 1.463 ± 0.349 μg/mL 2 and 6 hours after administration, respectively, and imatinib serum concentrations were 11.683 ± 1.086 μg/mL and 1.432 ± 0.353 μg/mL 2 and 6 hours after administration, respectively (Figure 9A-B). These results are consistent with those published by others that used comparable TKI doses in lean mice (59–62). Next, we quantified serum concentrations 2h after gefitinib and imatinib administration in obese mice after the first, fourth and eighth treatment. Gefitinib serum concentrations were 4.157 ± 0.546 μg/mL, 4.114 ± 0.108 μg/mL and 4.257 ± 0.165 μg/mL after the first, fourth and eighth administration, respectively, demonstrating how gefitinib administration reached steady-state concentrations after the first administration, and did not accumulate across the treatment duration (Figure 9C). The serum concentration of gefitinib in obese mice 2h after the first administration (4.157 ± 0.546 μg/mL) was comparable to the serum concentration of gefitinib achieved 2h after a single dose of gefitinib in lean mice (3.857 ± 0.263 μg/mL). In contrast, serum concentrations of imatinib in obese mice were 1.432 ± 0.353 μg/mL, 5.700 ± 1.404 μg/mL and 6.167 ± 1.308 μg/mL after the first, fourth and eighth administration, respectively, demonstrating that imatinib administration accumulates in the serum during the treatment regime (Figure 9D). The serum concentration of imatinib in obese mice 2h after the first administration (1.432 ± 0.353 μg/mL) was significantly lower than the serum concentration achieved 2h after a single dose of imatinib in lean mice (11.683 ± 1.086 μg/mL), suggesting that HFD-feeding affects drug metabolism of the TKI imatinib. By the eighth administration of TKI in obese mice, a comparable serum concentration of each TKI had been achieved (4.257 ± 0.165 μg/mL for gefitinib vs. 6.167 ± 1.308 μg/mL) despite the five-fold higher dose of imatinib that was used in our experiments.

**Figure 9.**
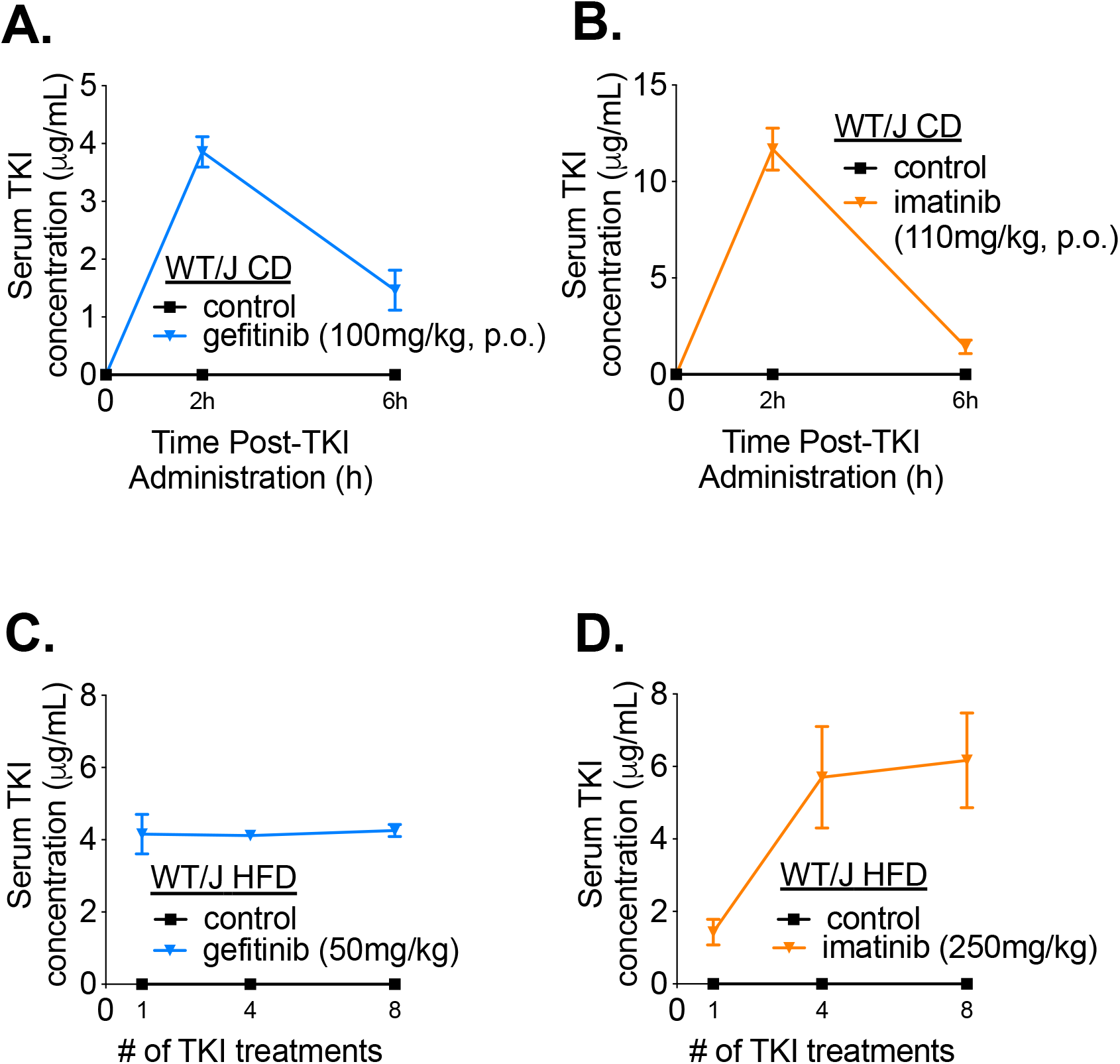
Quantification of gefitinib and imatinib in the serum of lean and obese mice. Quantification of TKI concentrations in mouse serum using liquid chromatography coupled to tandem mass spectrometry in lean WT/J mice fed a CD, at t=0, 2 and 6 h after a single administration of A) gefitinib (100mg/kg, p.o.), or B) imatinib (110mg/kg, p.o.), and in obese WT/J mice fed a HFD for 10 weeks before repeated administration of A) gefitinib (50mg/kg, p.o.), or B) imatinib (250mg/kg, p.o.). In obese mice, serum TKI levels were quantified in samples collected 2h after the 1^st^, 4^th^ and 8^th^ administration (see Figure 1). Values are mean± SEM. n = 1 for control group (not detected at any time point) and n=6-7 for TKI treatment groups.

## DISCUSSION

TKIs inhibit many on- and off-target kinases and this may underpin their divergent effects on blood glucose and insulin. The goal of this work was to understand how RIPK2 modifies the effects of certain TKIs on blood glucose and insulin in obese in mouse models. We previously demonstrated that the TKI gefitinib attenuates both NOD1-mediated dysglycemia in mice and increased adipocyte lipolysis, which are hallmarks of metabolic disease. We initially hypothesized that inhibition of the NOD1-RIPK2 signalling axis would contribute to the beneficial effect of the TKI gefitinib on glycemia during obesity. However, RIPK2 is also required to propagate downstream signalling from NOD2 and work from our lab has already shown that MDP is an insulin-sensitizing postbiotic that engages NOD2 (40,41). The cross-talk and magnitude of contributions of NOD1 versus NOD2 pathways in modifying glycemic control during obesity are not well understood. The potential value of targeting RIPK2 depends on the relative importance of NOD1 signalling compared to NOD2 signalling and their combined contributions to glycemic control and insulin sensitivity in metabolic disease states.

Gefitinib is known to inhibit RIPK2, but contrary to our initial hypothesis gefitinib treatment improved glucose control in all the obese mice that we tested, indicating that this TKI lowers blood glucose independent of NODs or RIPK2. Interestingly, lower insulin (coupled with lower blood glucose) during a glucose challenge was only observed in *Ripk2^-/-^* mice treated with gefitinib. This is an important stand-alone result since lowering of hyperinsulinemia is positioned to attenuate both insulin resistance and obesity and possibly even diabetic complications (63–65). Thus, these results suggest that deletion of RIPK2 improves the insulin sensitizing effect of gefitinib because glucose-lowering occurs concurrently with lower blood insulin only in *Ripk2^-/-^*mice treated with gefitinib. One possibility is that deletion of RIPK2 increases drug availability for other kinases and this warrants further investigation.

We next tested imatinib in obese HFD-fed mice, since this TKI has no reported inhibitory activity against RIPK2 *in vitro* (29). We previously demonstrated that imatinib does not inhibit glycemic or lipolytic consequences of NOD1-RIPK2 signalling (45). Here, we also showed that imatinib does not inhibit glycemic consequences of NOD2-RIPK2 signalling. Imatinib lowered obesity-induced glucose tolerance, but *increased* insulin secretion during glucose challenge. This data suggests that imatinib works through a mechanism distinct from gefitinib to improve glycemic control during obesity. Imatinib increases insulin secretion and reduces blood glucose levels whereas gefitinib lowers blood glucose levels without a concomitant increase in insulin. Augmented insulin secretion by imatinib is consistent with reports that inhibition of c-Abl, either with imatinib or c-Abl siRNA knockdown, enhances insulin secretion in beta cells *in vitro* (27). Furthermore, two case reports in cancer patients being treated with the TKIs dasatinib and sunitinib, which inhibit c-Abl to a similar or greater degree compared to imatinib, have been reported to increase serum c-peptide levels (a marker of insulin secretion) (9,14). It has been proposed that imatinib alleviates specific cellular-stress mechanisms that offer protection against beta-cell failure and prolongs or enhances the ability of the pancreas to secrete high amounts of insulin to counter insulin resistance at the level of the tissues. Beyond improved glycemic control, assessing a TKI’s individual effect on insulin is an important consideration when retasking TKIs as a therapeutic for obesity-induced insulin resistance or diabetic complications. Hyperinsulinemia itself has been implicated in the pathogenesis of insulin resistance and increasing evidence suggests high circulating insulin levels are a frontrunner in the development of obesity, driving subsequent metabolic dysfunction (66).

Additional molecular targets common to many TKIs have been investigated, such as PDGFRβ and c-Kit. In studies that investigated the glucose lowering effects of imatinib and sunitinib in models of Type 1 Diabetes, these effects were replicated by specific antibodies to PDGFRβ, but not c-Kit, suggesting that PDGFRβ inhibition may play a role in mediating glucose lowering effects, at least in animal models of Type 1 Diabetes (19,20). Interestingly, gefitinib does not have reported inhibitory activity against c-Kit or PDGFRβ (29). We found that imatinib increased insulin secretion to lower blood glucose in obese mice. Hence, investigating other kinase targets beyond PDGFRβ, which lowers glucose in Type 1 Diabetes models that do not have a large capacity to increase insulin secretion. These results also highlight the need to study the molecular targets of TKIs in models of Type 1 versus Type 2 diabetes. Treatment with gefitinib or imatinib did not result in widespread changes in gene expression of inflammatory- and ER stress-related markers in metabolic tissues. Based on the numerous targets of each TKI, it is plausible that different TKIs work through different mechanisms to exert divergent effects on insulin secretion versus improved insulin sensitivity in peripheral tissues to manifest effects on blood glucose regulation.

It is important that all our results using gefitinib or imatinib occurred despite no change in body mass or adiposity. We used male mice with obesity established after 10 weeks of HFD for experiments involving repeated treatment with TKIs. We confirmed that our selected doses did not impact food consumption, body mass, or adiposity in obese mice across the duration of the treatment period. It is possible that the doses used here would alter body mass in other mouse models. Despite the dose of imatinib required to improve blood glucose (250mg/kg) being 5-fold higher than the dose of gefitinib (50mg/kg), comparable serum concentrations were observed after the 8^th^ treatment (6.167 ± 1.308 μg/mL and 4.257 ± 0.165 μg/mL, respectively), at which point glucose control was assessed.

Here we demonstrate that NOD-RIPK2 signalling is not the defining molecular target of TKIs that underpins their effects on blood glucose. Multiple TKIs, including gefitinib and imatinib, lower blood glucose in obese mice, independent of RIPK2. However, RIPK2 participated in TKI-induced lowering of blood insulin in response to an oral glucose load. Thus, RIPK2 can discriminate the effects of some TKIs on insulin sectretion, but ultimately, the molecular target(s) that promote glucose-lowering remain elusive. Our data provides evidence that RIPK2 is a target that limits the insulin sensitizing potential of certain TKIs. Furthermore, we show different TKIs, including ones that do not inhibit RIPK2, such as imatinib, can improve blood glucose control by augmented insulin secretion (Figure 10). It is possible that other TKIs that work via the same molecular target(s) as gefitinib to lower blood glucose, but do not inhibit RIPK2, could be well-positioned to lower blood glucose *and* lower blood insulin. This concept warrants further investigation for re-tasking TKIs in diabetes treatment. Because each TKI has its own unique profile of selectivity and specificity, we propose individual TKIs being investigated for diabetes should be evaluated for their effects on blood glucose control and insulin dynamics, as well as their potential to inhibit RIPK2. Taken together, our results suggest that an in-depth exploration of how TKIs alter *both* insulin and glucose metabolism is necessary to efficiently harness the therapeutic potential of these drugs in obesity and diabetes.

**Figure 10.**
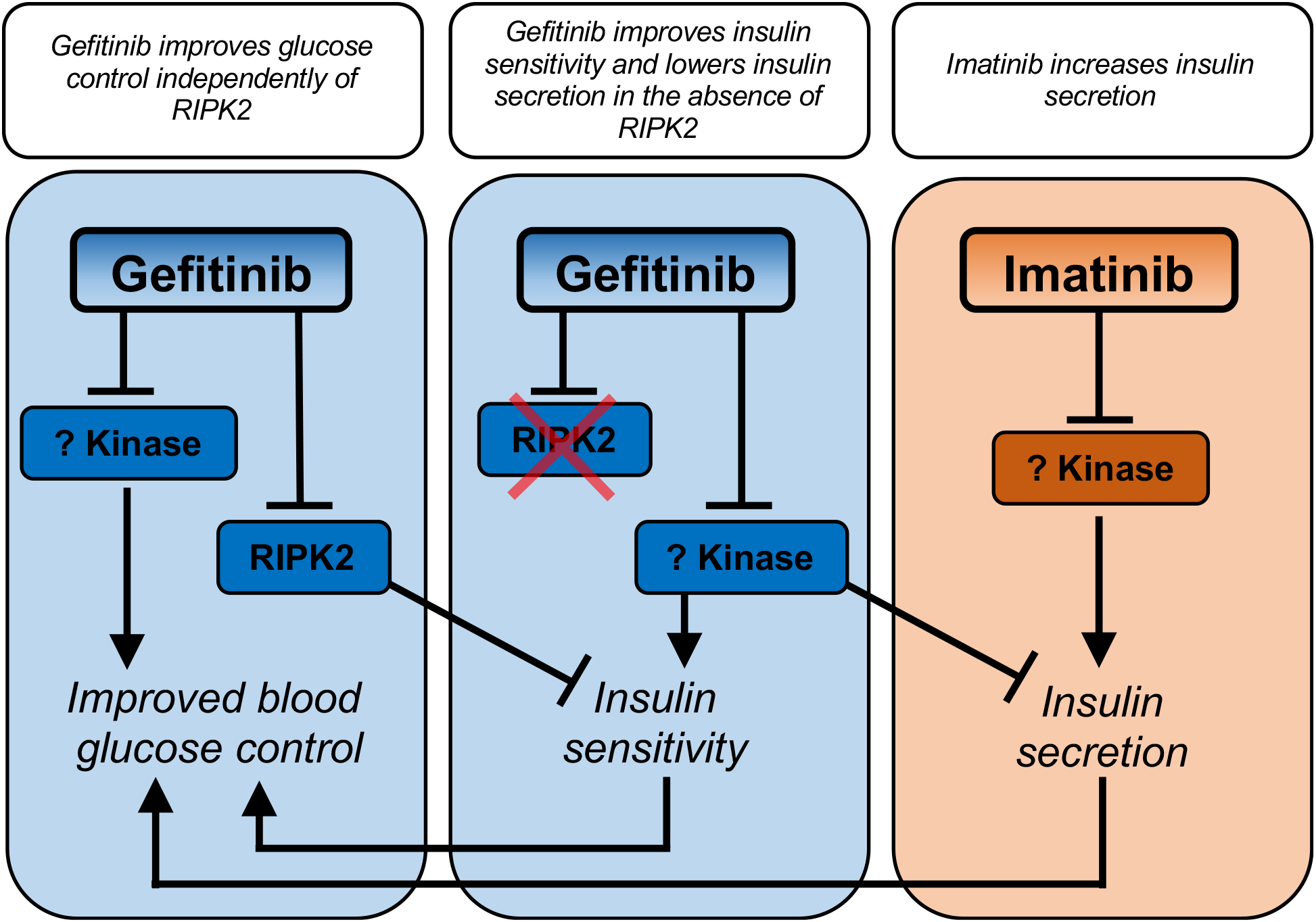
Inhibition of RIPK2 limits insulin sensitizing potential of a TKI. TKIs can operate independently of NOD1-RIPK2 or NOD2-RIPK2 signalling to improve blood glucose. TKIs, such as gefitinib, that inhibit RIPK2 signalling lower glucose stimulated insulin secretion only in obese *Ripk2^-/-^* mice, but not in wild type mice. RIPK2 inhibition limits the ability of gefitinib to simultaneously lower insulin and glucose and hence, the insulin sensitizing potential of this TKI. Other TKIs, such as imatinib increase insulin secretion to achieve lower blood glucose, independent of actions on RIPK2.

## Author Contributions

BMD, KPF, NGB, and JFC conducted experiments. BMD and JDS designed experiments, contributed to discussion, wrote and edited the paper. All authors have reviewed the manuscript.

## Abbreviations

TKIs: tyrosine kinase inhibitors
PRRs: pattern recognition receptors
TLRs: Toll-like receptors
NLRs: Nod-like receptors
MDP: muramyl dipeptide
PDGFRβ: platelet-derived growth factor receptor β
CD: control diet
HFD: high fat diet
GTT: glucose tolerance test
ITT: insulin tolerance test
OGSIS: oral glucose-stimulated insulin secretion

